# Postural constraints recruit shorter-timescale processes into the non-Gaussian cascade processes

**DOI:** 10.1101/2020.06.05.136895

**Authors:** Mariusz P. Furmanek, Madhur Mangalam, Damian G. Kelty-Stephen, Grzegorz Juras

**Affiliations:** Department of Physical Therapy, Movement and Rehabilitation Sciences, Northeastern University, Boston, MA 02115, USA; Institute of Sport Sciences, The Jerzy Kukuczka Academy of Physical Education, 40-065 Katowice, Poland; Department of Psychology, Grinnell College, Grinnell, IA 50112, USA

**Keywords:** center of pressure, inverse power law, postural control, quite stance, scale invariance

## Abstract

Healthy human postural sway exhibits strong intermittency, reflecting a richly interactive foundation of postural control. From a linear perspective, intermittent fluctuations might be interpreted as engagement and disengagement of complementary control processes at distinct timescales or from a nonlinear perspective, as cascade-like interactions across many timescales at once. The diverse control processes entailed by cascade-like multiplicative dynamics suggest specific non-Gaussian distributional properties at different timescales. Multiscale probability density function (PDF) analysis showed that when standing quietly while balancing a sand-filled tube with the two arms elicited non-Gaussianity profiles showing a negative-quadratic crossover between short and long timescales. A more stringent task of balancing a water-filled tube elicited simpler monotonic decreases in non-Gaussianity, that is, a positive-quadratic cancellation of the negative-quadratic crossover. Multiple known indices of postural sway governed the appearance or disappearance of the crossover. Finally, both tasks elicited lognormal distributions over progressively larger timescales. These results provide the first evidence that more stringent postural constraints recruit shorter-timescale processes into the non-Gaussian cascade processes, that indices of postural sway moderate this recruitment, and that more stringent postural constraints show stronger statistical hallmarks of cascade structure.

## 1. Introduction

Intermittency is a mathematical way to discuss subtle irregularities that characterize dexterous, goal-directed behavior [1,2]. Dexterous behavior hews close to task demands without becoming locked into those demands [3–5]. It requires mostly stable repertoires that can skitter beyond local task demands to meet longer-range contextual demands. For instance, comfortable walking uses the relatively stable cycle of swinging legs with a characteristic frequency [6,7], but at less local grains, walking a long distance outside in hot summer sun requires pausing for breath or detour through the shade. Posture is similarly intermittent. Postural dexterity entails swaying to support optic flow as well as eye and head movement [8,9]. Beyond satisfying local constraints of keeping the center of pressure (CoP) on the ground, the upright body sways to explore the surroundings—excessive sway topples the body, but insufficient sway leaves it poorly poised to enact or participate in ongoing events [10–12].

Intermittency reflects loose clustering of behavior around an average position as in an attractor basin and context-sensitive spreading-out into more extreme regions [13,14]. That is, whereas the bell-curve of a Gaussian distribution reflects a tendency to cling to the average position (e.g., hips over feet shoulder-width apart), intermittency entails long and heavy tails stretching outwards on either side of the average position (e.g., the way the torso may sway from side to side) [15,16]. The structure of probability density functions (PDFs) offers the possibility that we can evaluate just how rigidly behavior is locked into average position or how fluidly it can glide back and forth between meeting local task constraints and exploring the longer-scales of context beyond those local constraints. Specifically, the smoothness of the peak-to-tail transition offers the possibility that the ‘extremes’ of dexterous behavior are not random, disruptive interruptions. Instead, dexterous behavior is a much smoother, fluid cascade in which events at multiple scales are closely coordinated with one another. For instance, the extremely brief, small saccades of the eye cover targets made visible by a slower, larger turn of the head, and the turn of the head explores different relationships of overlapping visual surfaces as torso sway explores different regions of the available optic flow. Rather than uprooting stability and toppling the body suddenly from the two feet, the excursions of dexterous postural sway might grow relatively seamlessly from an overlapping set of physiological processes.

Cascade-like fluidity of intermittent sway can be quantified by examining PDFs of postural fluctuations. Whereas Gaussian-shaped curves have thin tails, thinner than those found in exponential distributions [17], the most fluid interactions amongst multiscaled processes should generate relatively heavier tails. A Gaussian shape follows from the independence among component processes converging on the average behavior [18]. In contrast, progressively smoother descents from peak to tail in a PDF of postural fluctuations would reflect the component processes’ multiplicative interaction, yielding a lognormal distribution [19]. Hence, we aim in this work to showcase analysis of postural fluctuations by examining intermittency as this non-Gaussian departure from a strictly short-tailed Gaussianshaped curve and into relatively lognormal PDF forms.

Analyses of intermittency abound in neuroscience, with multifractal detrended fluctuation analysis (MF-DFA) as the gold standard for demonstrating cascade-like dynamics [20,21] and recurrence quantification analysis [22] and entropy measures [23], each speaking to limited subsets of what MF-DFA encodes [24]. Nonetheless, one MF-DFA will not fit all because intermittency can and does slip from one statistical guise from another. The detrended variance may grow according to uniform power-law forms only for limited scales, suggesting change, instability, or disappearance of fractal patterns in the detrended variance [25]. Although interruptions in fractal-like temporal correlations could signal when overly strong task constraints overwhelm intermittency [21], such interruptions may also reflect the evolution of system manifesting intermittency in heavy PDF tails. Although Gaussian-defined variance serves admirably for minor deviations of PDFs from Gaussianity (e.g., exponential and gamma), estimates of temporal correlations like MF-DFA are progressively less likely to diagnose cascade-like intermittency when PDFs extend heavy tails (e.g., lognormal or even infinite-variance Lévy-stable PDFs) across extreme events [26,27].

PDF non-Gaussianity thus complements portraits of intermittency when multifractal-type methods falter [28]. Multiscale PDF analysis resolves these issues and makes an elegant companion to MF-DFA. Whereas fitting tails has been a standard practice using maximum likelihood estimation (MLE) [19], multiscale PDF overcomes two major limitations in this prior practice. First, although Gaussian distributions are clear indications of independence among constituent components that sum together to produce non-intermittent behavior, we cannot guarantee that another distribution shape will entail the opposite. For instance, when components interact like in a cascade rather than sum, those cascade-like interactions produce distributions with heavier tails, like, the lognormal distribution or the power-law distribution [19]. However, although heavy-tailed failures of Gaussian shapes are the consequence of cascade-like intermittency, the converse is not necessarily true. That is, heavy-tailed distributions can arise from non-cascade origins [29]. Second, PDFs are no less prone to crossovers and bends, or discontinuities in the PDF curve from one shape to another over a wide range of scales [30]. Whereas MF-DFA generates profiles of variance across multiple scales (measuring the distribution of residuals), multiscale PDF generates a PDF of displacements over multiple scales (measuring the distribution of displacements). Multiscale PDF analysis quantifies non-Gaussianity in intermittency through a metric 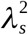 that increases as variance deviates from Gaussian form, with the expectation that cascade-like behavior would show a monotonic decrease from small to large scales [31,32]. 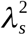 is not a simple cut off but estimates lognormal variance above and beyond the Gaussian variance [33].

Although this approach is itself blind to the structure across time, it has two benefits: 1) assessing PDF-based intermittency across scales; and 2) when we compare the results to those for linear phase-randomized surrogates, diagnosing precisely when non-Gaussian form arises from nonlinear interactions across time scale. Notably, we should expect crossovers no less than in standard PDF analysis, but using multiscale PDF in conjunction with MF-DFA and related intermittency metrics offers a new way to understand crossovers. Minimally, crossovers in 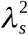 will indicate different scales constraining intermittency generating the PDF tail, and for now, we may ask whether these crossovers wax and wane as MF-DFA and similar measures become inversely worse or better at portraying intermittency. Crossovers can reflect different modes of control needed on either side of scale-dependent thresholds, e.g., in position or velocity [25]. Although such thresholds may be important to postural control in quiet standing, a destabilizing task constraint could relax this threshold and allow postural control to manifest a cascade process specific to the destabilizing task constraint.

We applied multiscale PDF analysis to study postural control in a tube-balancing task in which participants stood quietly and bimanually balanced sand- and water-filled tubes as stably as possible. Water is more fluid than sand and is thus more likely to destabilize posture than densely packed sand. Destabilizing task constraints prompt postural control to enrich its motor variability at shorter timescales, as stability is more fragile and needs to be reestablished more often, requiring sensory corrections more frequently or on much more subtle timescales. Hence, the groundswell of non-Gaussianity prompts the postural control to fan out a wider variety of displacements, allowing it to respond with much more than just a preponderance of the average-sized displacement. Therefore, we predicted that balancing the sand-filled tube would damp or elicit small-scale intermittency and elicit non-Gaussianity profiles showing a negative-quadratic crossover (i.e., an inverted U-shaped curve across timescales; Hypothesis-1a) and water-filled tube would cancel the crossover between short and long timescales (Hypothesis-1b). The dependent variable in both cases is the lambda relationship with log-timescale. These scale-dependent constraints on postural sway should not merely manifest in crossovers in non-Gaussianity; we expected that they should show a close association with indices of postural sway, including the temporal-correlation measures as from MF-DFA. We expected that indices of postural sway would moderate the quadratic form of the lambda curve, specifically entailing an interaction of postural-sway indices with the quadratic effect of log-timescale (Hypothesis-2).

## 2. Methods

### 2.1. Participants

Ten adult men (*mean±1SD* age = 21.4±1.1 years) with no neurological, muscular, or orthopedic conditions participated after providing institutionally-approved informed consent.

### 2.2. Experimental setup and procedure

We asked the participants to stand quietly with their feet shoulder-width apart and bimanually balance sand- and water-filled tubes (*l×d* = 150×20 cm, *m* = 8 kg) with as much stability as possible (Fig. 1). The experimenter ensured that the tubes didn’t touch any parts of the body except the hands. A force platform (100 Hz, AMTI Inc., Watertown, MA) recorded ground reaction forces for the entire 30 s duration. The participants completed five 30-s trials per task. The order of the trials was pseudo-randomized across participants.

**Fig. 1.**
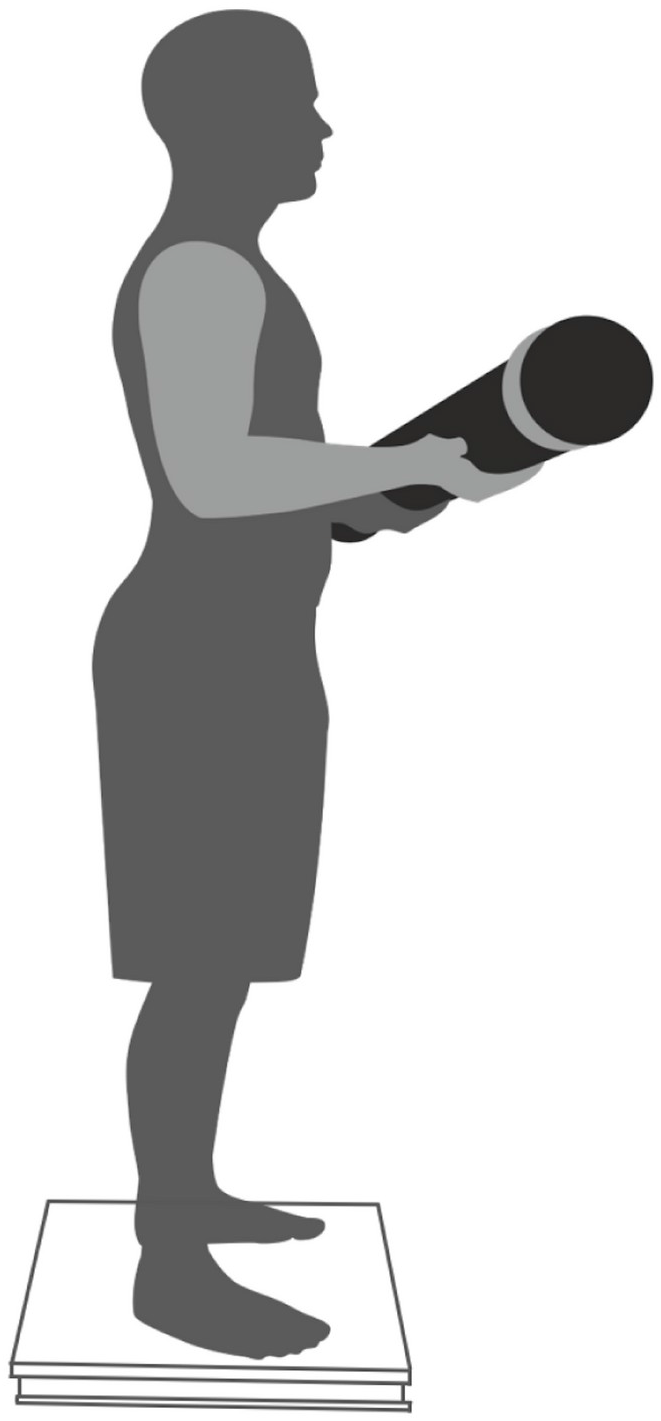
Experimental task. Each participant stood quietly while bimanually balancing a sandor water-filled tube for 30 s.

### 2.3. Data processing

All data processing was performed in MATLAB 2019b (Matlab Inc., Natick, MA). Trial-by-trial ground reaction forces yielded a 2D foot center of pressure (CoP) series, each dimension describing CoP position along anterior-posterior (AP) and medial-lateral (ML) axes. Each 30-s trial yielded 3000-sample 2D CoP series and 2999-sample 2D CoP displacement series. Finally, a 1D CoP planar Euclidean displacement (PED) series described postural sway along the transverse plane of the body. Iterated Amplitude Adjusted Fourier Transformation (IAAFT) provided phase-randomized surrogates using original series’ spectral amplitudes to preserve only linear temporal correlations [34].

### 2.4. Canonical indices of postural sway

Three linear indices were computed for each CoP PED series: (i) Mean of all fluctuations (*CoP_Mean*), (ii) Standard deviation (CoP_*SD*), and (iii) Root mean square (*CoP_RMSE*).

Four nonlinear indices were computed for each CoP PED series: (i) Sample entropy (*CoP_SampEn*), indexing the extent of complexity in the signal [35]. We used *m* = 2 and *r* = 0.2 for computing the *SampEn*. (ii) Multiscale sample entropy (CoP_*MSE*), indexing signal complexity over multiple timescales [35]. We used *m* = 2, *r* = 0.2, and *τ* = 20 for computing the *MSE*. (iii) We used detrended fluctuation analysis [36] to compute Hurst’s exponent, *H*_fGn_, indexing temporal correlations in original CoP PED series (CoP_ *H*_fGn_Original_) and randomly shuffled versions of each series (CoP_ *H*_fGn_Shuffled_) using the scaling region: 4, 8, 12,... 1024 [37], as well as the difference between the two (CoP_*H*_fGn_diff_OS_). Shuffling destroys the temporal structure of a signal. (iv) Multifractal spectrum width, Δ*α*, indexing the extent of multifractal temporal correlations in the signal. Chhabra and Jensen’s direct method [38] was used to compute Δ*α* for the original CoP series (CoP_Δ*α*_Original_) and the iterative amplitude adjusted Fourier transform (IAAFT) surrogate (CoP_Δ*α*_Surrogate_) [39], as well as the difference between the two (CoP_Δ*α*_diff_OS_).

### 2.5. Multiscale probability density function (PDF) analysis

Multiscale PDF analysis characterizes the distribution of abrupt changes in CoP PED series { *b*(*t*) } using the PDF tail. The first step is to generate { *B* (*t*) } by integrating { *b*(*t*) } after centering by the mean *b_ave_* (Fig. 2, top):

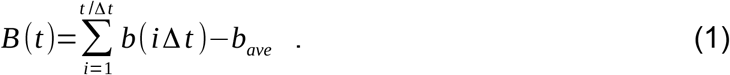

**Fig. 2.**
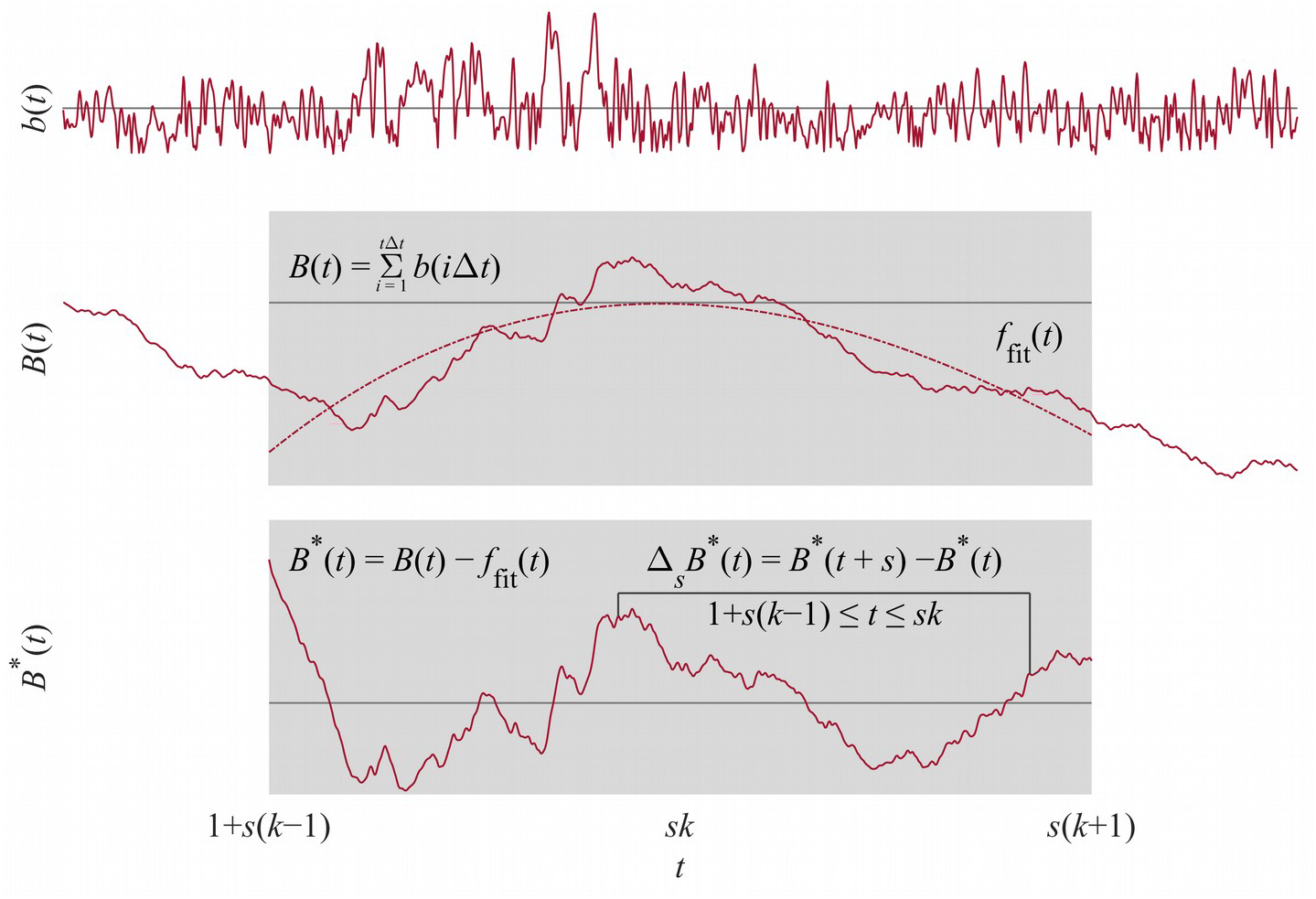
Schematic illustration of multiscale probability density function (PDF) analysis. See Section 2.5 for details.

A 3^rd^ order polynomial detrends { *B* (*t*) } within *k* overlapping windows of length 2*s*, *s* being the timescale (Fig. 2, middle). Intermittent deviation Δ *_s_B* (*t*) in *k^th^* window from 1+*s*(*k*–1) to *sk* in the detrended time series { *B^d^*(*t*)=*B* (*t*)–*f_fit_*(*t*) } is computed as Δ*_s_B^d^*(*t*)=*B^d^*(*t*+*s*)—*B^d^*(*t*), where 1 +*s*(*k* – 1)≤*t*≤*sk* and *f_fit_*(*t*) is the polynomial representing the local trend of { *B*(*t*) }, of which the elimination assures the zero-mean probability density function in the next step (Fig. 2, bottom). Finally, Δ *_s_B* is normalized by the SD (i.e., variance is set to one) to quantify the PDF.

To quantify the non-Gaussianity of Δ*_s_B* at timescale *s*, the standardized PDF constructed from all the Δ *_s_B* (*t*) values is approximated by the Castaing model [31], with *λ_s_* as a single parameter characterizing the non-Gaussianity of the PDF. *λ_s_* is estimated as

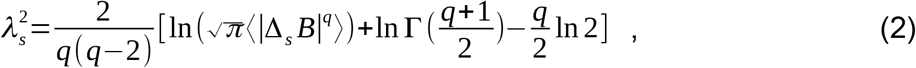

where 〈|Δ*_s_B*|*^q^*〉 denotes an estimated value of *q^th^* order absolute moment of { Δ*_s_B* }. As 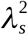 increases, the PDF becomes increasingly peaked and fat-tailed (Fig. 3, left). 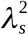 can be estimated by Eq. (2) based on *q^th^* order absolute moment of a time series independent of *q*. Estimating 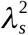 based on 0.2^th^ moment (*q* = 0.2) emphasizes the center part of the PDF, reducing the effects of extreme deviations due to heavy-tails and kurtosis. We used 0.2^th^ moment because estimates of 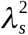 for a time series of ~ 3000 samples are more accurate at lower values of *q* [33].

**Fig. 3.**
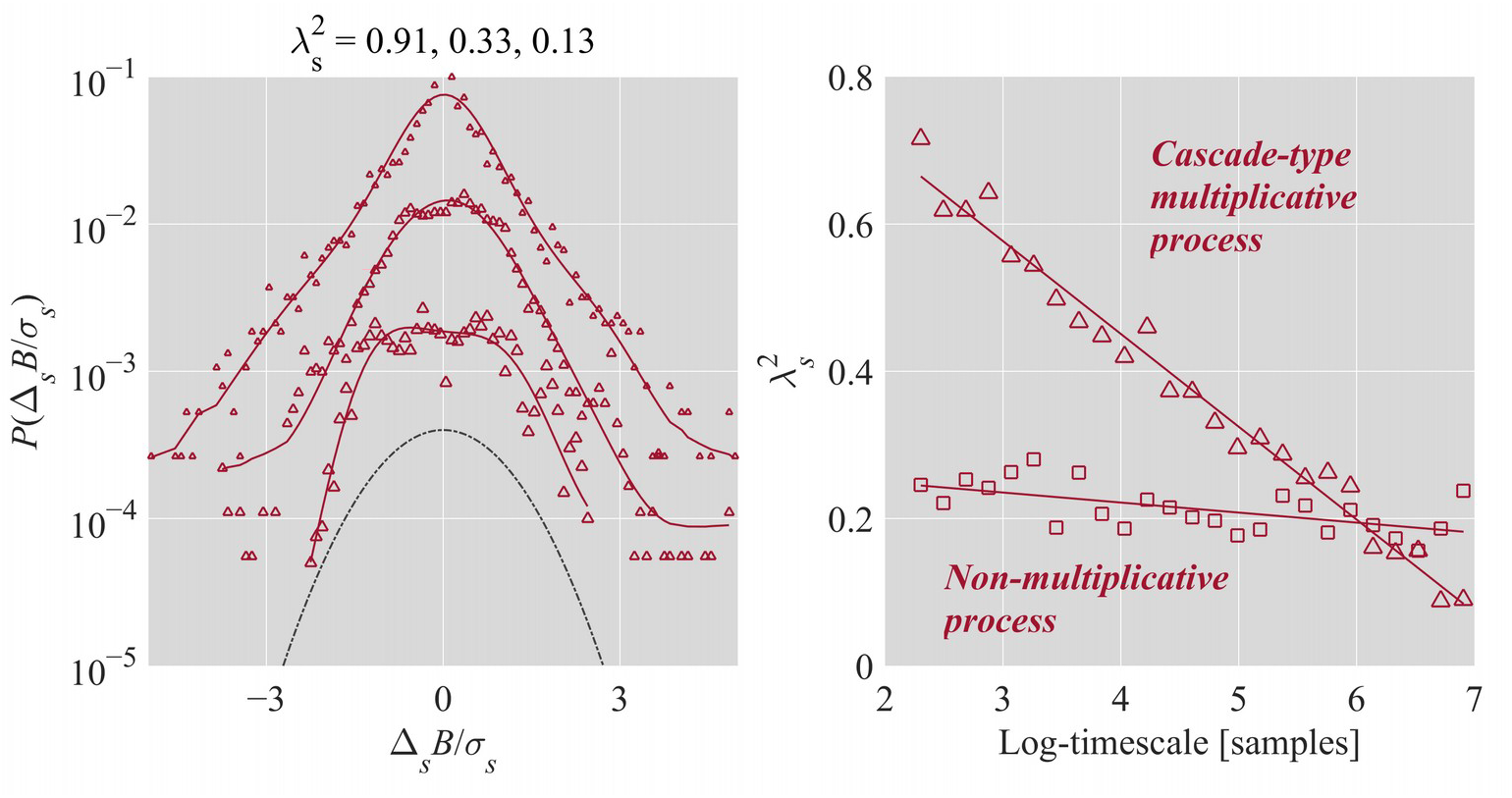
Schematic illustration of non-Gaussianity index *λ_s_*. Left: The relationship between 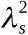 and shapes of PDF plotted in linear-log coordinates for 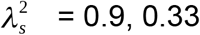, and 0.13 (from top to bottom). For clarity, the PDFs have been shifted vertically. As 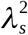 increases, the PDF becomes increasingly peaked and fat-tailed. As *λ_s_* decreases, the PDF increasingly resembles the Gaussian (dashed line), assuming a perfect Gaussian as 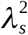 approaches 0. Right: Cascade-type multiplicative processes yield the inverse relationship 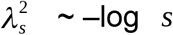.

Cascade-type multiplicative processes yield the inverse relationship 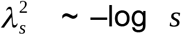 (Fig. 3, right) [32]. For the present purposes, we quantified 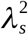 for each original CoP PED series and corresponding IAAFT surrogate at timescales of 5 to 1000 samples (i.e., 50 ms to 10 s) at steps of 5 samples (50 ms).

Multiscale PDF analysis of non-Gaussianity and traditional MLE methodology using Akaike Information Criterion (AIC) differ in their emphasis. In contrast to traditional MLE, which is disproportionately sensitive to the less-populated tails, multiscale PDF analysis addresses the central, better-populated bulk of the PDF. Hence, we also included MLE to complement multiscale PDF analysis, specifically to portray the analytical effect of sensitivity to different portions of the distribution. We used AIC to test which among a power law, lognormal, exponential, or gamma distribution characterized the PDF at each timescale.

### 2.6. Statistical analysis

To test for crossovers in non-Gaussianity, a linear mixed-effects (LME) model using *lmer* function [40] in R package *Ime4* tested 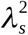 vs. log-timescale curves for orthogonal linear and quadratic polynomials, for interactions with grouping variables (Task × Original, where Original encoded differences in original series from surrogates) and with indices of postural sway (Section 2.4.). Statistical significance was assumed at the alpha level of 0.05 using R package *lmerTest* [41]. To test how lognormality changed with log-timescale, a generalized linear mixed-effects (GLME) model fit changes in Lognormality as a dichotomous variable (Lognormal = 1 vs. Exponential = 0) using orthogonal linear, quadratic, and cubic polynomials and tested interaction effects of grouping variables (Task × Original) with those polynomials using glmer [42] in *lme4*. We report convergence within 670 function evaluations, well below the 10000 default for glmer(). The model output returned no convergence errors and very small (< 0.001) maximum absolute relative Hessian-scaled gradients.

## 3. Results

Figure 4 provide a schematic of multiscale PDF characterization of postural sway in representative participants balancing sand- and water-filled tubes for 30 s. The following text focuses on the higher-order interactions’ coefficients and omits detail on lower-order control effects that only appear to ensure proper specification of interactions [43].

**Fig. 4.**
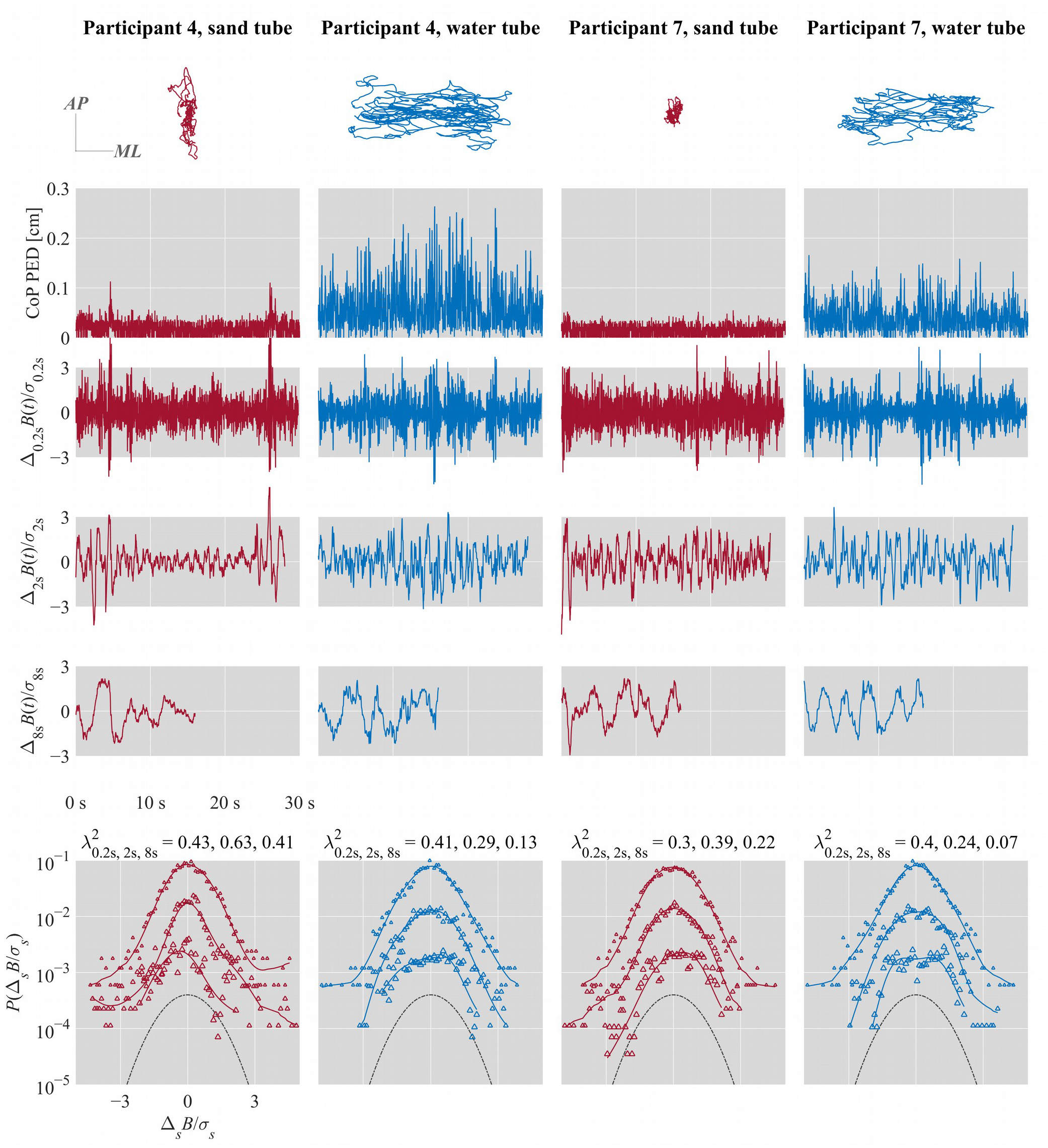
Multiscale PDF characterization of postural sway in representative participants balancing sand- and water-filled tubes for 30 s. From top to bottom: CoP trajectories along the anterior-posterior (*AP*) and medial-lateral (*ML*) axes. CoP PED series. {Δ*_s_ B*(*i*) } for *s* = 0.2, 2, and 8 s. Standardized PDFs (in logarithmic scale) of { Δ*_s_B*(*i*) } for *s* = 0.2, 2, and 8 s (from top to bottom), where *σ_s_* denotes the *SD* of { Δ*_s_B*(*i*) }.

### 3.1. Effects of task on 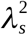 vs. log-timescale curves

In the sand-filled condition, 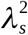 vs. log-timescale curves showed stronger linear decrease (*B* = –3.30, *P* < 0.001) and stronger negative quadratic, downward-facing parabolic form (*B* = –2.50, *P* < 0.001) for the original CoP PED series than the surrogates (Table S1; Fig. 5, left). Balancing the water-filled tube elicited a significant reversal of this parabolic form (*B* = 0.50, *P* = 0.014). Specifically, in the water-filled condition, 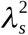 vs. log-timescale curves for the original CoP PED series showed both a stronger linear decrease (*B* = –2.93, *P* < 0.001).

**Fig. 5.**
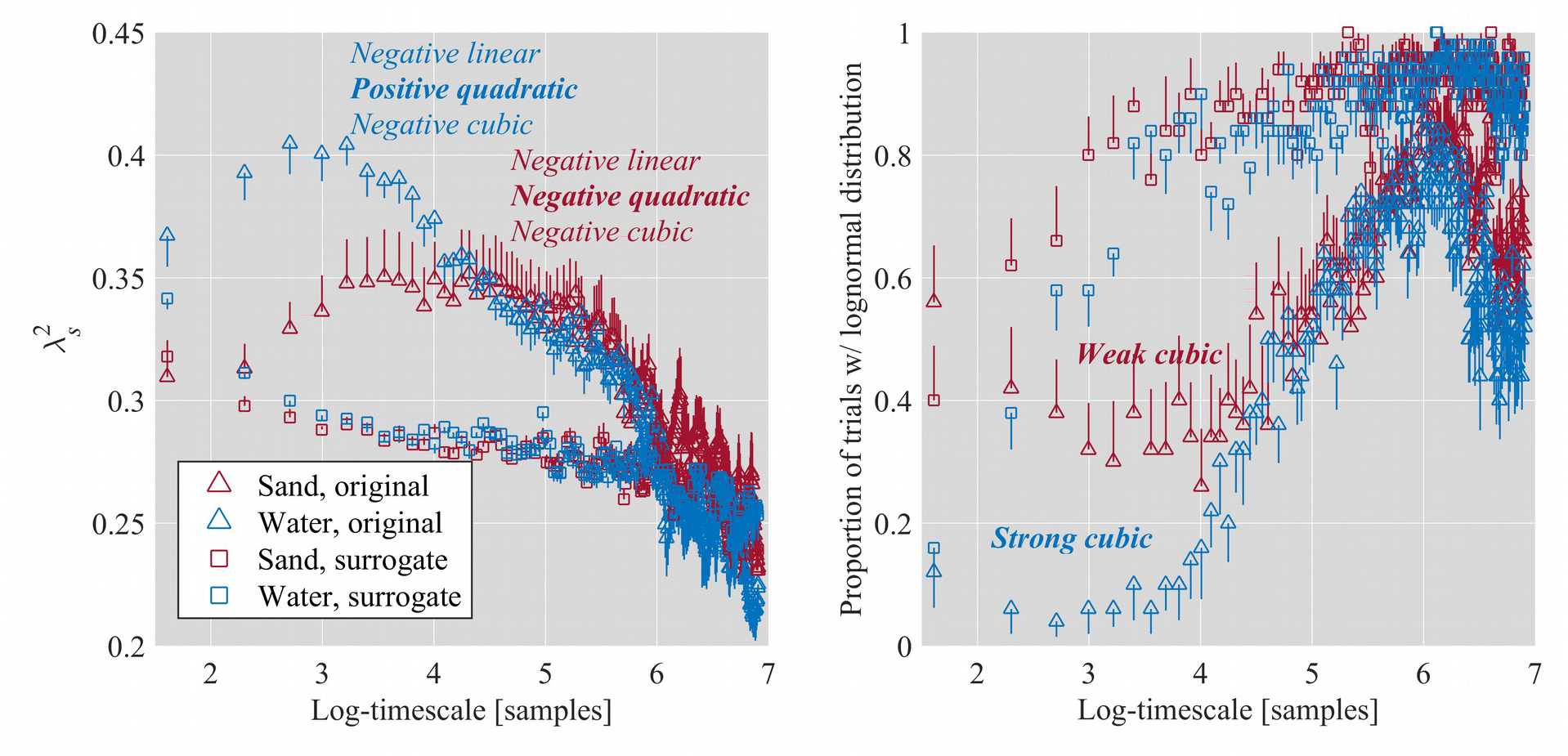
Log-timescale dependence of mean 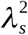 (left; Section 3.1; Tables S1 and S2) and empirical proportion of trials with a lognormal distribution (right; Section 3.2; Table S3), while balancing sand- and water-filled tubes for 30 s. Vertical bars indicate ±1*SEM* (*N* = 10).

Adding interactions of postural-sway indices with the quadratic relationships significantly improved model fit, *χ*^2^(99) = 3994.04, *P* < 0.0001, and showed that the only significant change in the parabolic form of 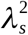 vs. log-timescale curves depended on the indices of postural sway. That is, the main effect of quadratic relationships between 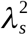 and log-timescale, the difference in this quadratic form between the original CoP PED series vs. the surrogates, and the difference in this quadratic form due to the water-filled tube for the original COP PED series yielded non-significant coefficients (*P*s > 0.05; Table S2).

Otherwise, indices of postural sway moderated the quadratic form of 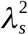 vs. log-timescale curves (Table S2) directly addressing the strength of the downward-opening parabolic form. Increases in CoP_*RMSE* accentuated the negative quadratic form in the original COP PED series (*B* = –1.37×10^6^ *P* = 0.032). Increases in CoP_*Mean*, CoP_*SD*, and CoP_*MSE* accentuated the negative quadratic form for the original COP PED series vs. the surrogates in the water-filled condition (*B* = 1.06×10^6^ *P* = 0.037; *B* = 8.04×10^5^, *P* < 0.038; and *B* = −6.66, *P* = 0.009; respectively). Other effects reversed the negative quadratic form in the sand-filled condition. Increases in CoP_*Mean*, CoP_*SD*, CoP_*MSE*, and CoP_Δ*α*_diff_OS_ contributed positive quadratic coefficients for the original COP PED series vs. the surrogates (*B* = 1.09×10^6^, *P* = 0.032; *B* = 8.28×10^5^, *P* = 0.033; *B* = 14.49, *P* < 0.001; and *B* = 2.73×10^3^, *P* < 0.001, respectively). Increases in CoP_*H*_fGn_ contributed quadratic coefficients in the water-filled condition (*B* = 15.35, *P* = 0.028). Increases in CoP_*RMSE* contributed positive quadratic coefficients for the original COP PED series vs. the surrogates in the water-filled condition (*B* = 1.33×10^6^, *P* = 0.037).

### 3.2. Effects of task on the growth of lognormality in postural sway with log-timescale

Fig. 5, right shows the proportion of PDFs for which MLE returns lowest AIC for lognormal fit. All other MLE results indicated exponential distributions. In the original CoP PED series, lognormality in postural sway increased as a cubic function of log-timescale, with significant positive linear (*B* = 19.09, *P* < 0.001) and significant negative cubic forms (*B* = - 71.87, *P* < 0.001; Table S3). The quadratic component of the lognormality vs. log-timescale relationship was not significant (*p* > 0.05). In contrast, in the surrogates, the growth of lognormality showed a parabolic form with a significant linear increase and quadratic decrease with log-timescale in the sand-filled condition (*B* = 53.07, *P* < 0.001; and *B* = - 34.32, *P* < 0.001, respectively). These linear and quadratic effects became stronger in the water-filled condition (*B* = 20.43, *P* < 0.001; and *B* = −40.34, *P* < 0.001, respectively), but crucially, this difference between the sand- and water-filled conditions was the same for the original CoP PED series and the surrogates.

## 4. Discussion

Multiscale probability density function (PDF) analysis supported Hypothesis-1a: balancing the sand-filled tube elicited quiet standing with non-Gaussianity profiles showing a negative-quadratic crossover between short and long timescales. The analysis also supported Hypothesis-1b: balancing the water-filled tube elicited simpler monotonic decreases in non-Gaussianity in the form of a positive-quadratic cancellation of the negative-quadratic crossover, suggesting the recruitment of shorter-timescale processes into the non-Gaussian cascade processes. Indices of postural sway governed the appearance or disappearance of the crossover, thus supporting Hypothesis-2. Finally, both tasks elicited heavy tails over progressively larger timescales, confirming the cascade structure of postural control and a strong sensitivity of multiscale PDF analysis to non-Gaussianity in sway, occurring alongside MLE evidence of lognormal distributions. These results provide the first evidence that stringent postural constraints recruit shorter-timescale processes into the non-Gaussian cascade processes, that indices of postural sway moderate this recruitment, and that more stringent postural constraints show stronger statistical hallmarks of cascade structure, strengthening the evidence of cascade processes in postural control.

A noteworthy finding is that postural sway reflects definite postural adjustments amenable to task constraints. Higher potential of heavy-tailed fluctuations at sufficiently longer timescales, as indicated by the change in PDF from exponential to lognormal at these longer timescales, indicate definite postural adjustments. Balancing the water-filled tube was associated with both higher non-Gaussianity and potential of heavy-tailed fluctuations at shorter timescales than balancing the sand-filled tube, indicating that definite postural adjustments can be selectively recruited at shorter timescales.

Non-Gaussianity confirmed by a new analytical method is a major step forward for modeling the cascade-type intermittency of postural sway. The cascade structure of postural control is no airy mathematical abstraction unmoored from the physiological details. Instead, we can pinpoint the physiological substance: the vast network of connective tissues and the extracellular matrix that has been imagined as a multifractal tensegrity in which the components hang together under tensional and compressional forces at multiple scales [44]. This work has not tested tensegrity-themed hypotheses about non-local effects across the body, like other studies have tested [45]. Nonetheless, it is essential to note that although previous work on such hypotheses acknowledges tensegrity-free, specifically neural-based explanation, no known alternatives to the tensegrity hypothesis predicts intermittency arising from multifractally formed nonlinear interactions across scales.

Invoking tensegrity may seem threatening to neuroscientific perspectives, but it should not. Certainly, multifractal geometry aligns as well with neuronal avalanches [46] found in the nervous system, but again, to our knowledge, the tensegrity proposal is alone in offering a physiological basis for predicting intermittency arising from multifractally formed nonlinear interactions across scales. In contrast, current perspectives on the nervous system’s development and function emphasize networks of tension as the large-scale context framing the electrical and molecular dynamics of the brain [47]. Indeed, Ingber [48] articulates that this large-scale tensional context provided by the extracellular matrix is a necessary precondition for the typically developed, sensory-specific action potential in the peripheral nervous system. Meanwhile, Turvey and Fonseca [44] review the facts that, despite some differences in the brain ECM as compared to the more peripheral ECM, there are clear avenues through which bodywide fascial tissue could interact with the fascia of the brain (e.g., through myodural lines potentially reaching from the body to the meninges and acting on astrocyte networks).

It can be no small coincidence that tensegrity and neuronal-avalanche models all converge on the same multifractal geometry [48] that inspired the development of multiscale PDF analysis. Indeed, estimates of multifractality along this physiological cascade well predict the flow of perceptual information through the body and the coordination of disparate body parts [49,50]. Further, we see direct evidence that multifractal signatures of nonlinearity across time through DFA have direct relationships to cascade structure as estimated through multiscale PDF analysis, suggesting convergence of evidence through different analytical sources. Now, multiscale PDF appears as a crucial complement to more popular multifractal analyses when intermittency evolves to generate heavy-tailed PDFs. This wider view of intermittency will equip future research to clarify the view on how cascade-like intermittency might change with task and with age in providing support for action.

## Supplementary material

**Table S1**. Coefficients of the linear mixed-effects (LME) model examining the effects of task on 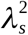 vs. log-timescale curves.

**Table S2**. Coefficients of the LME model examining the effects of task on 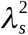 vs. log-timescale curves for orthogonal linear and quadratic polynomials, for interactions with indices of postural sway.

**Table S3**. Coefficients of the generalized linear mixed-effects (GLME) model examining the effects of task on the growth of lognormality in postural sway with log-timescale.

## Author contributions

M.P.F., M.M., and D.G.K-S. conceived and designed research; M.P.F. performed experiments; M.M. and D.G.K-S. analyzed data; M.P.F., M.M, and D.G.K-S. interpreted results of experiments; M.M. prepared figures; M.M. and D.G.K-S. drafted manuscript; M.P.F., M.M., D.G.K-S., and G.J. edited and revised manuscript; M.P.F., M.M., D.G.K-S., and G.J. approved final version of manuscript.

